# Potential role of heat shock protein 90 in regulation of hyperthermia-driven differentiation of neural stem cells

**DOI:** 10.1101/598441

**Authors:** Lei Wang, Yi Zhuo, Zhengwen He, Ying Xia, Ming Lu

**Author notes:** Corresponding authors: Ying Xia; Ming Lu. Equal contributors:Lei Wang, Yi Zhuo.

## Abstract

**Objective:** Our previous studies indicated that hyperthermia may play a role in differentiation of neural stem cells and that hypoxia inducible factor-1(HIF-1) was critical in this process. Heat shock protein 90 (Hsp90) is one of the most common heat-related proteins and involved in HIF-1 expression by regulating its activity and stabilization. Here, we hypothesized that HSP90 may be involved in regulation of hyperthermia-driven differentiation of neural stem cells(NSCs). We also investigated whether the HSP90 activity exert its regulatory action via HIF-1 pathway and the transcriptional level of the target genes of HIF-1.

**Method:** The cultured NSCs were divided into three groups: an hyperthermic treatment group(40NSC) which NSCs was induced under 40°C temperature; a control group(37NSC) which NSCs was induced under 37°C temperature; an hyperthermic treatment and HSP90-inhibited group(40NSC+GA) which NSCs was induced with 0.5μM HSP90 inhibitor Geldanamycin(GA) under 40°C temperature. We examined cells HSPa and HIF-1a expression during a time window of 5 days(12h, 1d, 3d, 5d) post-differentiation. The expression HSPα, HIF-1α, VEGF (vascular endothelial growth factor) and erythmpoietin(EPO) of during a time window was evaluated by RT-qPCR. The proportion of Tuj-1 positive differentiated cells were observed by flow cytometry.

**Result:** Hyperthermia promoted neuronal differentiation of NSC, and this effect could be blocked by HSP90 inhibitor GA. We observed the up-regulation of HSP90 during hyperthermia treatment, and that the protein levels of HIF-1α changed depending of the GA treatment. GA could not inhibited HSP90α expression but suppressed HSP activity and decreased the expression HIF-1α protein. Inhibition of HIF-1α expression by GA could consequently affect expression of its targeted genes such as VEGF and EPO.

**Conclusion:** Hyperthermia promote differentiation of NSCs into neurons. HSP90 involved in the regulation of hyperthermia-driven differentiation of NSC, and the mechanism is related to HIF-1α and its downstream gene activation.

## Introduction

Neural stem cells(NSCs), which have the potential of highly self-renewal and can be differentiate into other types of neural tissue cells including neurons, astrocytes and oligodendrocytes under certain micro-environment[1-3]. Based on these biological characteristics of NSCs, NSCs transplantation bring new hope of treatment for central nervous system(CNS) injury and neurodegenerative disorders. However, most of transplanted cells are surrounded by a hostile microenvironment that does not support neuronal differentiation, the proportion of neuron cells differentiated from NSCs remains at a low level[4-6]. Thus, how to induce NSCs to differentiate into neural cells with a higher proportion is recently one of the cutting-edge fields in neuroscience research[7-8]. Fate determination and differentiation of NSCs is a complex physiological procedure that dependent on intrinsic and extrinsic cues, such as growth factors, transcription factors and signaling pathways activation[9-11].

Until recently, the effects of temperature on the differentiation of NSCs have been largely ignored. Our previous study demonstrated that hyperthermia conditioning could stimulate olfactory ensheating cells(OECs) over-expressed hypoxia inducible factor 1α(HIF-1α). Hyperthermic preconditioned OECs could induced NSCs differentiate into neuron more efficiently, and the mechanism involved in the HIF-1 signaling pathway[12]. In addition, hyperthermia promotes neuronal and glial fate specification in NSCs in the early stages of differentiation[13]. However, the mechanism of hyperthermia regulating differentiation of NSCs is still not clear.

Heat shock protein 90 (Hsp90) is one of the most common heat-related proteins and involved in HIF-1 expression by regulating its activity and stabilization. Various stress factors such as hyperthermia, hypoxia and viral infections can induce the expression of HSP90. We supposed that HSP90 might be involved in regulation of hyperthermia-driven differentiation in NSCs. A Research have indicated that hyperthermia-driven neural differentiation is accompanied with change of expression pattern of HSP90[14]. HSP90 is an essential protein that controls the activity and stabilization of HIF-1α[15], HSP90 could directly interact with the PAS domain of HIF-1α to make it keep stabilization[16].

Therefore, in this study, we hypothesized that HSP90 may be involved in the regulation of hyperthermia-driven differentiation of NSCs. In addition, we also investigated whether the HSP90 activity exert its regulatory action via HIF-1 pathway and the transcriptional level of its target genes.

## Materials and methods

### NSCs culture and induction of neural differentiation

Three-day newborn Sprague-Dawley(SD) rats were used. The ethical committee of Central South University approved all experiments involving rats and protocol.

Cells derived from SD rat cortex and hippocampus tissue were mechanically dissociated. The single cell suspension was cultured in DMEM/F12 medium(Gibco) supplemented with 2% B27(Abcam), 20 ng/ml EGF(Abcam) and 20 ng/ml bFGF(Abcam). The cells were incubated at 37°C and 5% CO2 and full humidity[12].

### Group of experiment

Cells were induced in DMEM/F12 medium contains 1% fetal bovine serum(FBS,Hyclone). The induced NSCs were divided into three groups. 40NSC: An hyperthermic treatment group which NSCs was induced with 1%FBS under 40°C; 37NSC: a control group which NSCs was induced with 1%FBS under 37°C; 40NSC+GA: a hyperthermic treatment and HSP90-inhibited group(40NSC+GA) which NSCs was induced with 1%FBS and 0.5μM HSP90 inhibitor Geldanamycin(GA, Cayman) under 40°C.

### Real-time qPCR

Cultured cells washed thrice with PBS and treated with TRIzol(Invitrogen) reagent and reverse-transcripted for cDNA synthesis with SuperScript III cDNA synthesis kit. Each cDNA subpopulation was subjected to polymerase chain reaction amplification using the specific primers. The sense and antisense primers for each marker were as follows: HSP90α, F: GAAATTGCCCAGTTAATGTCC, R: ATTCCAATGCCAGTATCCAC; HIF-1α, F: CTCATCAGTTGCCACTTCC, R: CCTAGAAGTTTCCTCACACGTA; VEGF, F: CTGGACCCTGGCTTTACTGCT, R: ACACCGCATTAGGGGCACAC; EPO, F: ACCCTGCTGCTTTTACTATCCTT, R: TACCTCTCCAGAACGCGACT; The PCR products were mixed with a loading buffer (0.25% bromophenol blue, 0.25% xylene cyanol, and 40% sucrose) and separated on 2% agarose gels. The data was analyzed using MxPro QPCR software.

### Western blot

All differentiated cells lysates cantaining 50 μg proteins were subjected to gel electrophoresis. And then all protein were transferred to PVDF membranes. The blots were blocked in 4% BSA in TBST solution for 30 min at room temperature and then incubated at 4-°C overnight with the primary antibody: HSP90α(Proteintech, 1:1000), HIF-1α(CST, 1:1000), β-actin(Proteintech, 1:5000). After incubation with secondary antibodies at room temperature for 1 h, the blot was visualized using Chemi Doc XRS imaging system (Bio-Rad)

### Immunofluorescence and flow cytometry

All differentiated cells were wished with PBS three times and fixed by alcohol. Cells were incubated with the primary antibody overnight at 4°C. The following primary antibodies were used: rabbit anti-Tuj-1 (Abcam, 1:1000) for neurons. And the cells were incubated with fluorochromecon jugated secondary antibodies for 1 h at room temperature. Cells were plated into test tubes at a density of 1×10^5^/mL. Cell fluorescence signals were determined immediately using flow cytometry with a FACS Caliber instrument (Becton Dickinson). The analysis was performed using Cell Quest Software (Becton Dickinson).

### Statistical analysis

All values are expressed as mean ± SEM. Statistical comparisons were performed in SPSS 16.0. Student’s two-tailed *t* test was used for comparing experimental groups, and a P value *<*0.05 was considered significant.

## Result

The primary NSCs were abtained as described in “Material and methods”. NSCs rapidly proliferated in the medium, forming small neurospheres after 5 days (Fig.1A). Neurospheres were positive for the NSCs marker Nestin(Fig 1B).

**Fig. 1.**
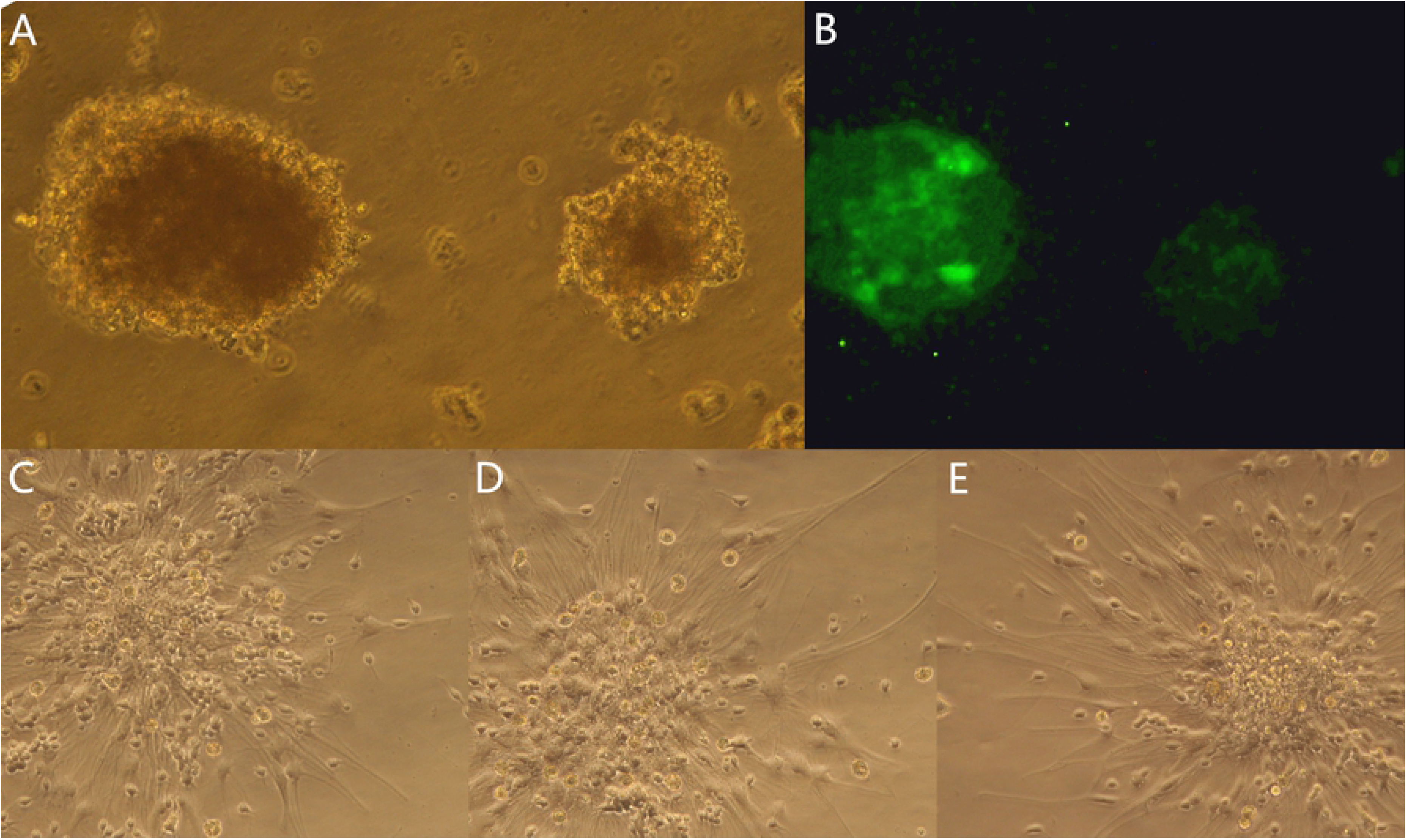
Characterization and differentiation of NSCs. A: phase-contrast image of NSCs globes cultured 5d in NSCs culture medium. B: Immunostaining of NSCs with Nestin antibody. C-F: Morphological changes in 37NSC, 40NSC+GA and 40NSC group after 5-day differentiation. Most differentiated cells in the 37NSC group(C) and 40NSC+GA group(D) were small, rounded, and triangular with two or three processes, while cells in the 40NSC group(E) were large, flat, and had an elongated shape with longer and wider processes.

For differentiation, the Neurospheres were cultivated in 1%-FBS-DMEM/F12 medium exposed to normal temperature(37°C, 37NSC) or hyperthermia(40°C, 40NSC) or hyperthermia with addition of GA(0.5μM, 40NSC+GA) for 5 days. After 5 days induction, differentiated cells in three groups exhibited neuronal or glia-like morphology. Differentiated cells in the 40NSC group exhibited large, flat, and had an elongated shape with longer and wider processes(Fig.1 E), while most cells in the 37NSC group and 40NSC+GA group were small, rounded, and triangular shape with two or three processes(Fig.1 C, D). To investigate influence of HSP90 on neural differentiation, cells at different time point of neural differentiation were monitored by flow cytometry.

Flow cytometry analysis revealed that the majority of differentiated cells in 40NSC group were positive for Tuj-1,the percentage of Tuj-1-positive cells in 40NSC group(51.91±3.69%) was significantly higher than that in 37NSC(31.46±2.21%) and 40NSC+GA groups(25.03±2.56%) after 3 days induction(Fig.2). The percentage of Tuj-1 positive cells in 40NSC(64.24±3.16%) was also significantly higher than that in 37NSC(35.43±2.65%) and 40NSC+GA groups(28.55±2.84%) after 5 days induction(Fig.2). The result of flow cytometry indicated that hyperthermia promoted neuronal differentiation of NSC, and this effect could be blocked by HSP90 inhibitor GA.

**Fig. 2.**
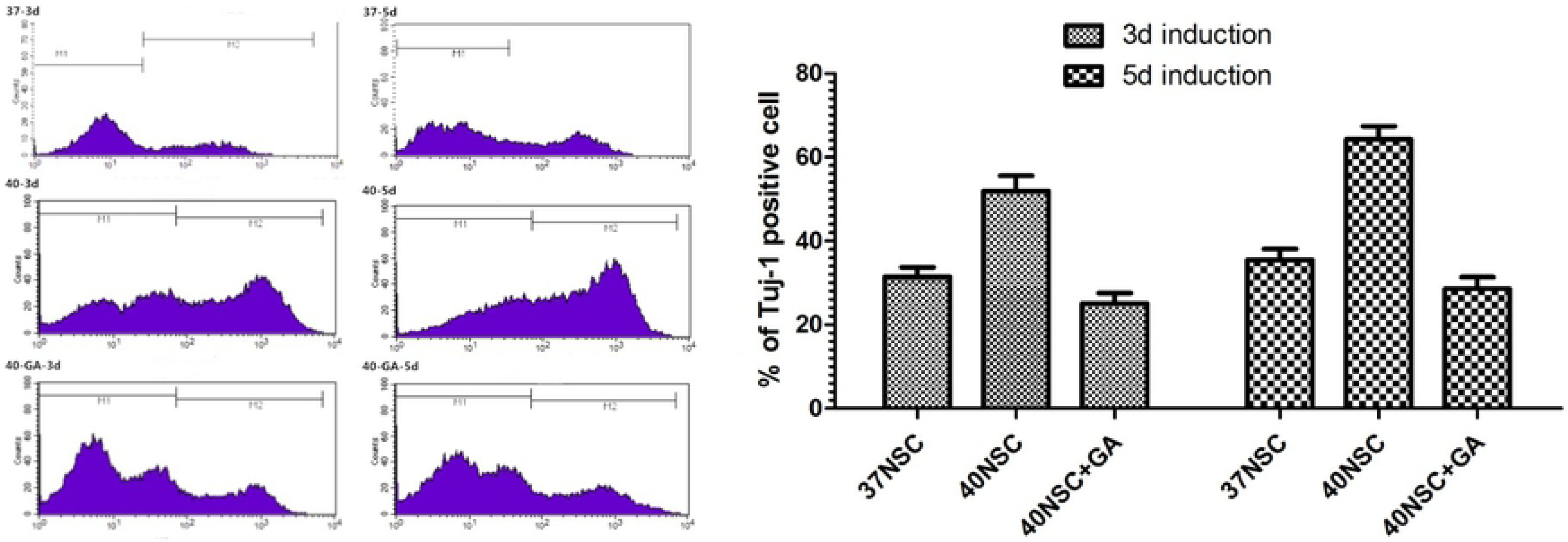
Flow cytometric analysis of Tuj-1-positive cells after 3 or 5 days induction.

To investigate the molecular mechanism in NSCs under hypoxia response, we examined the expression of HIF-1α and HSP90 in three groups. Cell lysates in three groups were extracted and tested by Western blot for HIF-1α and HSP90 expression. Figure 3 showed the expression of HSP90α levels increased in NSCs upon exposure to hyperthermia(both in 40 and 40NSC+GA groups) after 12h, 24h, 3d and 5d induction. However, no obvious statistical difference in HSP90 protein expression levels were detected between the two groups.

**Fig. 3.**
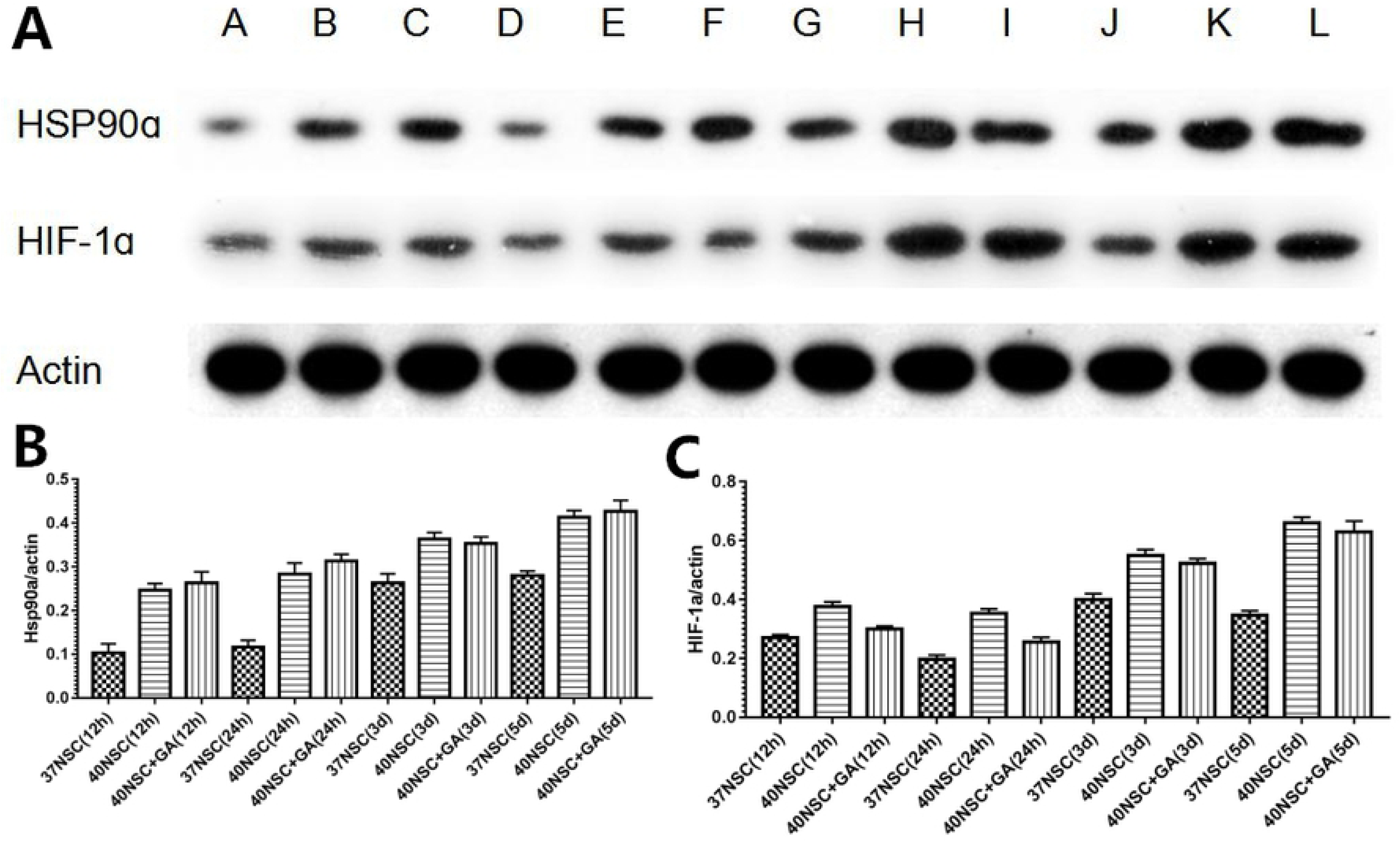
Western blot analysis of HSP90α and HIF-1α of induced NSCs(12h,24h,3d and 5d)·. A: 37NSC (12h induction); B: 40NSC (12h induction); C: 40NSC+GA (12h induction); D: 37NSC (24h induction); E: 40NSC (24h induction); F: 40NSC+GA (24h induction); G: 37NSC (3d induction); H: 40NSC (3d induction); I: 40NSC+GA (3d induction); J: 37NSC (5d induction); K: 40NSC (5d induction); L: 40NSC+GA (5d induction).

However, The HIF-1α protein in 40NSC group was significantly higher than that in the 37NSC group and 40NSC+GC group after 12h and 24h induction(Fig.3 A, C) and the band of HIF-1α expression became weaker during prolonged hyperthermia. As shown in Fig. 3, we observed the up-regulation of HSP90 during hyperthermia treatment, and that the protein levels of HIF-1α changed depending of the HSP90 inhibitor GA treatment. Hyperthermia-induced expression of HIF-1α was inhibited by GA as compared with its control. The result is similar to those that have been well documented[15]. HSP90 inhibitor GA could not inhibited HSP90α expression but suppressed HSP activity and decreased the expression HIF-1α protein.

To investigate the potential mechanism of HSP90α in NSC fate specification, we examined the mRNA level of HSP90α, HIF-1α, VEGF and EPO. VEGF and EPO were both regulated by HIF-1α, the levels of VEGF and EPO was evaluated to further examine whether GA inhibited the transcriptional control of HIF-1 to its targeted genes. RT-PCR showed that the mRNA level of HSP90α significant increased in NSCs exposed to hyperthermia environment(40NSC and 40NSC+GA groups) from 12h to 5d post-differentiation(Fig.4 A). HIF-1α was more abundantly expressed in the 40NSC than that in the 37NSC and 40NSC+GA group after induction(Fig.4 B). Expression of VEGF and EPO in 40NSC was also significant higher than those in 37NSC group and 40NSC+GA group(Fig.4 C, D). Hence, inhibition of HIF-1α expression by GA could consequently affect expression of its targeted genes such as VEGF and EPO.

**Fig. 4.**
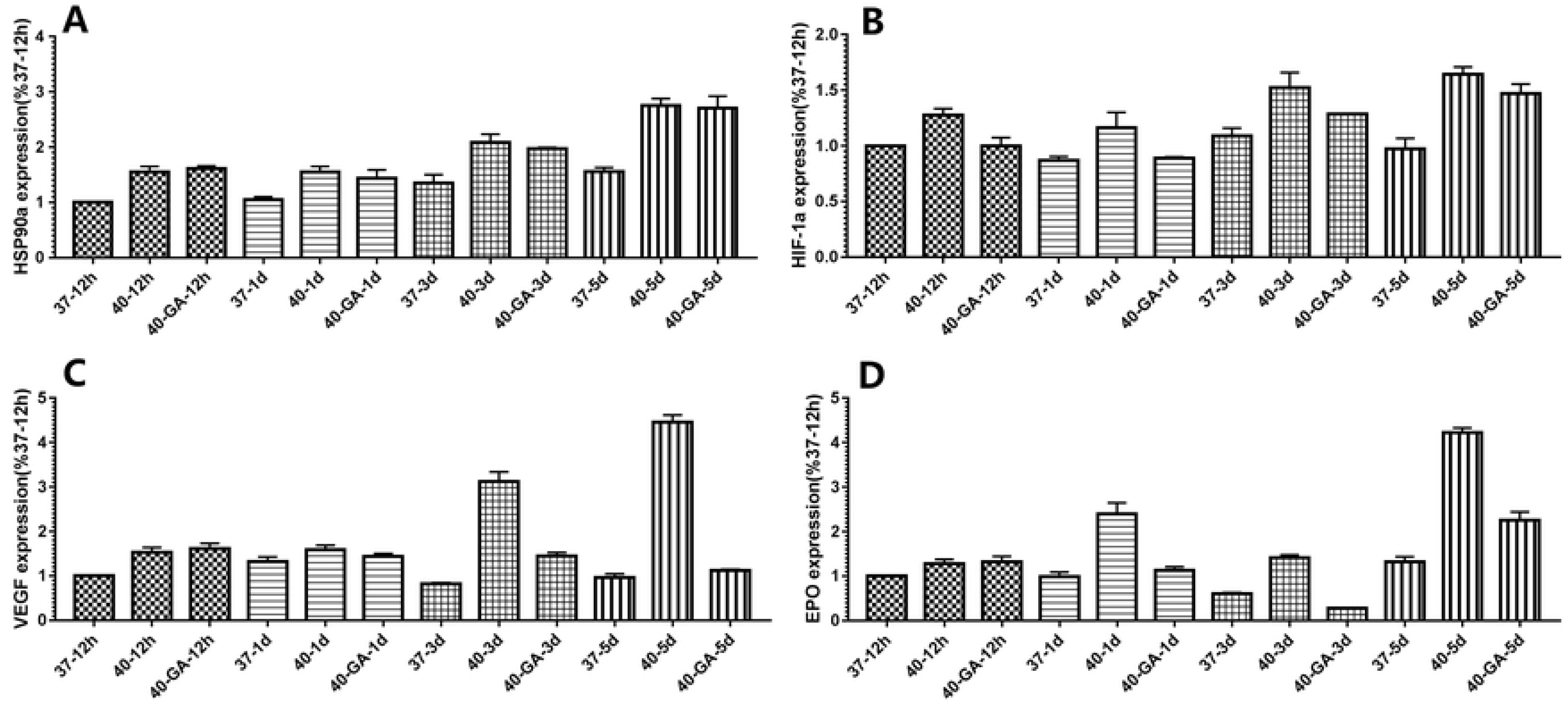
RT-qPCR analysis of HSP90α, HIF-1α, VEGF, EPO expression form 12h to 5d after differentiation.

## Discussion

Our date indicated that hyperthermia promoted neuronal differentiation of NSC, and this effect could be blocked by GA. HSP90α was critical in hyperthermia-driven differentiation of NSC. Hyperthermia condition could promote differentiation of NSCs into neurons with HSP90α over-expression, while this phenomenon could be specifically suppressed by GA. Although mRNA and protein expression of HSP90α was detected strongly in NSCs under hyperthermia surrounding whether treated with GA or not, both mRNA and protein expression levels of HIF-1α were significant decreased after treatment with GA under hyperthermia. This suggests that GA could significantly influence the expression of HIF-1α through regulation of HSP90 activity.

The above date indicated that the high expression of HSP90α induced by hyperthermia may be one important factors that promote differentiation of NSCs into neurons. HIF-1 is a known as downstream target of HSP90, it has been reported that HSP90α could make HIF-1α keep stabilization via interaction with PAS domain of HIF-1α[17]. In this study, our data indicated that the HSP90 inhibitor GA could not inhibited HSP90α expression but suppressed HSP activity and decreased the expression HIF-1α protein. HIF-1 translocation and activation in the nucleus results in the production of several downstream genes, such as vascular VEGF and EPO. The expression of EPO and VEGF, which were mainly regulated by HIF-1, was also reduced by GA.

HIF-1 composed of an α subunit and a β subunit, and the regulation of the HIF-1 activity depends mostly upon the α subunit[18]. HIF-1 is not only a low-oxygen sensor response to hypoxia but also is important regulator in NSCs. Previous studies reported that HIF-1α involved in the regulation of hyperthermia-driven catecholaminergic and dopaminergic differentiation of central nervous system stem cell, and the mechanism is related to HIF-1α and its downstream gene activation[19-20]. Knock out HIF-1α caused neural precursor cells impairment of survival and proliferation. The number of dopaminergic neuron differentiated of HIF-1α knockout mNPCs was also markedly decreased[21]. Further, over-expression of HIF-1α could stimulate NSC proliferation by activating the Wnt/β-catenin signaling pathway[22]. Increase in HIF-1α and VEGF expression promoted the proliferation and differentiation of NPCs and functional recovery following cerebral ischemia[23-24]. HIF-1α plays a central role not only in the regulation of NSPC response to hypoxia, metabolism and maintenance of the vascular environment of the neural stem cell niche[25], also is a regulator for fate decisions in the neural lineage[26]. Our earlier studies have shown that hyperthermia-conditioned olfactory ensheathing cells could induce NSCs differentiation into neuron more efficiently by the upregulation of HIF-1α and binding activity[12].

Previous study has shown that the endothelial cells promote NSCs proliferation and differentiation associated with VEGF, possibly by activating the Notch and Pten pathways[27]. Shifts in the VEGF isoforms result in transcriptome changes correlated with early NSCs proliferation and differentiation in mouse forebrain[28]. The VEGF signaling is a crucial target of Cav-1, Cav-1 can inhibit neuronal differentiation via down-regulations of VEGF[29]. In vivo experiment, Recombinant VEGF-NSCs transplantation following SCI is more efficacious compared to normal NSC transplantation[30]. All of these studies suggest that there is an essential linkage between VEGF and NSC differentiation.

Endogenous Epo/EpoR signaling contributes to neural cell proliferation in central nervous system in regions associated with neurogenesis[31]. Recombinant EPO significantly increased Akt activity and Ngn1 mRNA levels in neural progenitor cells, which was coincident with increases of neural progenitor cell proliferation, differentiation, and neurite outgrowth[32]. Recently a study report that EPO signaling promotes both neurogenesis and oligodendrogenesis following spinal cord injury[33].

Our date showed that the mRNA levels of HSP90α had no changes exposed to hyperthermia whether treated with GA or not, while the mRNA levels of HIF-1α, VEGF and EPO remarkably increased under hyperthermia compared to normal temperature. After treatment with GA under hyperthermia, the levels of HIF-1α, VEGF and EPO mRNA significant decreased. Hence, inhibition of HIF-1α expression by GA could consequently affect expression of its targeted genes such as VEGF and EPO.

HSP90 could regulate cell differentiation and proliferation under both physiological and pathological conditions[34]. HSP90 have a neuronal localization at all stages of postnatal development, which is tightly regulated by the cell cycle at neurulation. Some studies showed that HSP90 is necessary for cell differentiation and in cells starting to undergo neural morphogenesis[35]. HSP90 localized in cytoplasm and neurites during cell differentiation and increased about 6-fold in differentiated neural cells[34]. Further, HSP90 is involved in proliferation of embryonic neural stem/progenitor cells under hypoxia by regulating HIF-1α stabilization[15]. The HSP90 was made up of two isoforms, HSP90α (inducible form/major form) and HSP90β (constitutive form/minor form)[36]. Both mRNA and protein expression levels of HSP90β were lower than HSP90α, indicating that HSP90α might play a major role in growth of neural stem/progenitor cells[15].

The above date indicated that hyperthermia promote differentiation of NSCs into neurons. HSP90 involved in the regulation of hyperthermia-driven differentiation of NSC, and the mechanism is related to HIF-1α and its downstream gene activation.

## Conclusion

Hyperthermia promote differentiation of NSCs into neurons. HSP90 involved in the regulation of hyperthermia-driven differentiation of NSC, and the mechanism is related to HIF-1α and its downstream gene activation.

## Availability of data and materials

All data generated and/or analyzed during this study are included in this published article.

## List of Abbreviation

HIF-1: Hypoxia inducible factor-1
HSP90: Heat shock protein 90
NSCs: Neural stem cells
VEGF: Vascular endothelial growth factor
EPO: Erythmpoietin
DMEM/F12: Dulbecco’s modified Eagle’s medium and Ham’s
GA: Geldanamycin

